# Altering lipid droplet homeostasis affects *Coxiella burnetii* intracellular growth

**DOI:** 10.1101/112300

**Authors:** Minal Mulye, Brianne Zapata, Stacey D. Gilk

## Abstract

*Coxiella burnetii* is an obligate intracellular bacterial pathogen and a causative agent of culture-negative endocarditis. While *C. burnetii* initially infects alveolar macrophages, it has also been found in lipid droplet (LD)-containing foamy macrophages in the cardiac valves of endocarditis patients. In addition, transcriptional studies of *C. burnetii*-infected macrophages reported differential regulation of the LD coat protein-encoding gene perilipin 2 *(plin-2)*. To further investigate the relationship between LDs and *C. burnetii*, we compared LD numbers using fluorescence microscopy in mock-infected and *C. burnetii*-infected alveolar macrophages. On average, *C. burnetii*-infected macrophages contained twice as many LDs as mock-infected macrophages. LD numbers increased as early as 24 hours post-infection, an effect reversed by blocking *C. burnetii* protein synthesis. The observed LD accumulation was dependent on the *C. burnetii* Type 4B Secretion System (T4BSS), a major virulence factor that manipulates host cellular processes by secreting bacterial effector proteins into the host cell cytoplasm. To determine the importance of LDs during *C. burnetii* infection, we manipulated LD homeostasis and assessed *C. burnetii* intracellular growth. Surprisingly, blocking LD formation with the pharmacological inhibitors triacsin C or T863, or knocking out acyl-CoA transferase-1 (acat-1) in alveolar macrophages, increased *C. burnetii* growth at least 2-fold. Conversely, preventing LD lipolysis by inhibiting adipose triglyceride lipase (ATGL) with atglistatin almost completely blocked bacterial growth, suggesting LD breakdown is essential for *C. burnetii.* Together these data suggest that maintenance of LD homeostasis, possibly via the *C. burnetii* T4BSS, is critical for bacterial growth.

**IMPORTANCE:** Host neutral lipid storage organelles known as lipid droplets (LDs) serve as a source of energy, nutrients, and signaling lipids. LDs are associated with infection of the intracellular bacterial pathogen *Coxiella burnetii*, a significant cause of culture-negative endocarditis. While *C. burnetii* was found in LD-rich foamy macrophages in endocarditis patients, little is known about the host LD-*C. burnetii* relationship. We demonstrated *C. burnetii* Type 4B Secretion System (T4BSS)-dependent LD accumulation in macrophages, suggesting a T4BSS-mediated regulation of host LD homeostasis. Further, manipulating LD homeostasis significantly affected bacterial growth, indicating LDs play an important role during *C. burnetii* infection. As *C. burnetii* endocarditis has a 19% mortality rate even in treated patients, exploring the LD-*C. burnetii* association might identify novel therapeutic targets.

## INTRODUCTION

Lipid droplets (LDs) are dynamic cytoplasmic organelles which store cellular lipids in eukaryotic cells. LDs are uniquely comprised of a phospholipid monolayer surrounding a hydrophobic core of neutral lipids, primarily sterol esters and triacylglycerols (TAGs). LD assembly begins with neutral lipid synthesis, where fatty acyl CoA synthetases generate long chain fatty acids which are converted to sterol esters and triacyglycerols by acyl-CoA:cholesterol acyltransferase (ACAT) and acyl-CoA:diacylglycerol acyltransferase (DGAT), respectively. Progressive accumulation of neutral lipids in the ER leads to budding of the lipid ester globule surrounded by the ER membrane cytoplasmic leaflet, thus forming LDs (1, 2). Inversely, adipose triglyceride lipase (ATGL) (3) and hormone sensitive lipase (HSL) (4) mediate LD breakdown and release of free cholesterol and fatty acids. Functionally, LDs serve as intracellular lipid reservoirs for membrane synthesis or energy metabolism. In addition, LDs are linked to a range of cellular functions including protein storage, protein degradation and signaling (2, 5).

LDs are emerging as important players during host-pathogen interactions. During infection of host cells, Hepatitis C virus (HCV) (6) and Dengue virus (7) co-opt LDs as platforms for viral assembly and replication. Even though blocking LD formation attenuates HCV and Dengue virus replication *in vitro*, the importance of LDs during viral infection still remains elusive (8). LD numbers increased in host cells during infection with several pathogens including HCV (6) and Dengue virus (7), as well as the protozoan parasites *Trypanosoma cruzi* (9), *Plasmodium berghei* (10), *Toxoplasma gondii* (11), *Leishmania amazonensis* (12) and *Leishmania major* (13). In addition, infection with the intracellular bacterial pathogens *Chlamydia* spp. (14), *Mycobacterium* spp. (15-18), *Orientia tsutugamushi* (19), and *Salmonella typhimurium* (20) also led to increased LD numbers. Since fatty acids released from host TAGs and sterols can serve as carbon sources during infection (21), *C. trachomatis* (14, 22) and *M. tuberculosis* (15) are proposed to use LD components as a major source of energy and nutrients. Furthermore, in cells infected with *M. leprae* (23), *M. bovis* (16), *T. cruzi* (24), and *Leishmania infantum chagasi* (25), LDs serve as a source of prostaglandin and leukotriene eicosanoids, important signaling lipids which modulate inflammation and the immune response. These LD-derived eicosanoids potentially favor intracellular pathogen survival by downregulating the immune response (26).

LDs have been implicated during infection by *Coxiella burnetii*, a gram-negative intracellular bacterium and the causative agent of human Q fever. Primarily spread through aerosols, *C. burnetii* acute infection is characterized by a debilitating flu-like illness, while chronic disease results in endocarditis. Although *in vitro* and *in vivo C. burnetii* can infect a wide range of cells including epithelial cells and fibroblasts, the bacterium first infects alveolar macrophages during natural infection. Inside the host cell, *C. burnetii* directs formation of a specialized lysosome-like compartment called the parasitophorous vacuole (PV) which is essential for *C. burnetii* survival. PV biogenesis requires the *C. burnetii* type 4B secretion system (T4BSS), which secretes effector proteins into the host cell cytoplasm where they manipulate a wide range of cellular processes. While not established to be T4BSS-dependent, *C. burnetii* is thought to manipulate LDs and other components of host cell lipid metabolism (27-30). *C. burnetii*-containing LD-filled foam cells were found in heart valves of an infected patient (31), and LDs were observed in the *C. burnetii* PV lumen of infected human alveolar macrophages (32). Further, two separate microarray analyses reported differential regulation of the LD coat protein *plin-2* in *C. burnetii*-infected human macrophage-like cells (THP-1) (29, 30), suggesting *C. burnetii* induced changes in host cell LDs. Intriguingly, siRNA depletion of the phospholipase involved in LD breakdown, PNPLA2 (also known as ATGL), increased the number of *C. burnetii* PVs in HeLa epithelial cells (33). In addition, treatment of monkey kidney epithelial cells (Vero cells) with a broad spectrum antiviral molecule ST699 which localizes to host cell LDs inhibited *C. burnetii* intracellular growth (34). Despite these observations, the importance of LDs during *C. burnetii* infection is not known. In this study, we further examined the relationship between host LDs and *C. burnetii*. We observed a T4BSS-dependent increase in LD numbers in infected mouse alveolar and human monocyte-derived macrophage cell lines. Furthermore, manipulation of LD homeostasis significantly altered *C. burnetii* intracellular growth, thus strongly indicating that LDs play an important role during *C. burnetii* infection.

## RESULTS

### LD-associated genes are upregulated in *C. burnetii-*infected alveolar macrophages

To examine the role of LDs in *C. burnetii* pathogenesis, we first analyzed the expression of LD homeostasis-related genes in infected cells. As *C. burnetii* preferentially infects alveolar macrophages during natural infection, we utilized a mouse alveolar macrophage cell line (MH-S) previously established as a model for *C. burnetii* infection (35). Compared to mock-infected cells at day 1 post-infection, *C. burnetii* infection upregulated the expression of genes *acat-1*, *acat-2* and *fabp-4* by more than 1.2 fold (Figure 1). ACATs (acyl coA transferases) catalyze the formation of sterol esters, while FABP-4 (fatty acid binding protein-4) facilitates fatty acid transport for storage in LDs (36). Upregulation of LD-associated genes suggests LDs may increase during *C. burnetii* infection.

**Figure 1:**
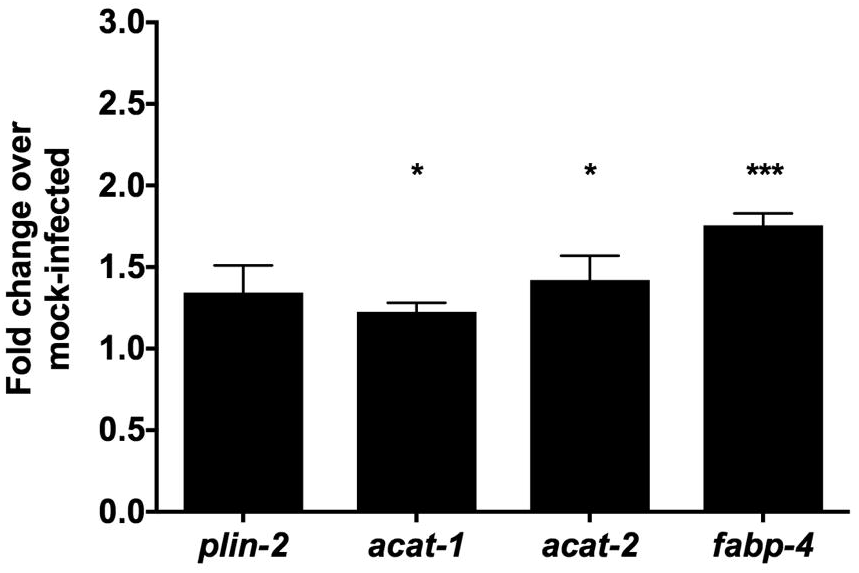
LO-associated genes are upregulated in *C. burnetii-infected* alveolar macrophages. MH-S macrophages were infected with wild-type C. *burnetii* and RNA was collected at day 1 post-infection. Gene expression analysis was performed using Applied Biosystems lipid regulated TaqMan Array. Fold change was calculated compared to mock-infected samples. Error bars show the mean of 6 independent experiments+/- SEM * =p<0.05, *** =p <0.001 compared to respective mock-infected cells as determined by two-tailed paired t-test.

### *C. burnetii* infection results in bacterial protein synthesis-dependent host cell LD accumulation

Previously, at 36 and 72 hours post-infection two separate microarray analyses of *C. burnetii*-infected THP-1 cells reported upregulation of the LD coat protein-encoding gene *plin-2* (29, 30). Mahapatra et al. also reported upregulation of *fabp-4* (29). Together with our gene expression analysis, this suggests LD formation may be altered during *C. burnetii* infection. To test this possibility, we stained LDs for the coat protein PLIN2 and quantitated LD numbers per cell in *C. burnetii*-infected cells by fluorescence microscopy. On an average, mock-infected had less than 50 LDs per cell during a 4 day experiment, irrespective of the time point (Figure 2A). In contrast, we observed a significant increase in LDs in infected cells, with an average of more than 90 LDs/cell at 1, 2 and 4 days after *C. burnetii* infection (Figure 2A, gray circles). Notably, LD accumulation occurred as early as 1 day post-infection, when the PV has not expanded and the bacteria are not in log growth (Figure 2D-E). To determine the role of *C. burnetii* in host cell LD accumulation, we blocked bacterial protein synthesis with chloramphenicol (29) and quantitated LD numbers at indicated times. Chloramphenicol-treated *C. burnetii*-infected cells contained 50 or less LDs per cell similar to mock-infected cells (Figure 2B and G). This result suggests *C. burnetii* protein synthesis is required for LD accumulation.

**Figure 2:**
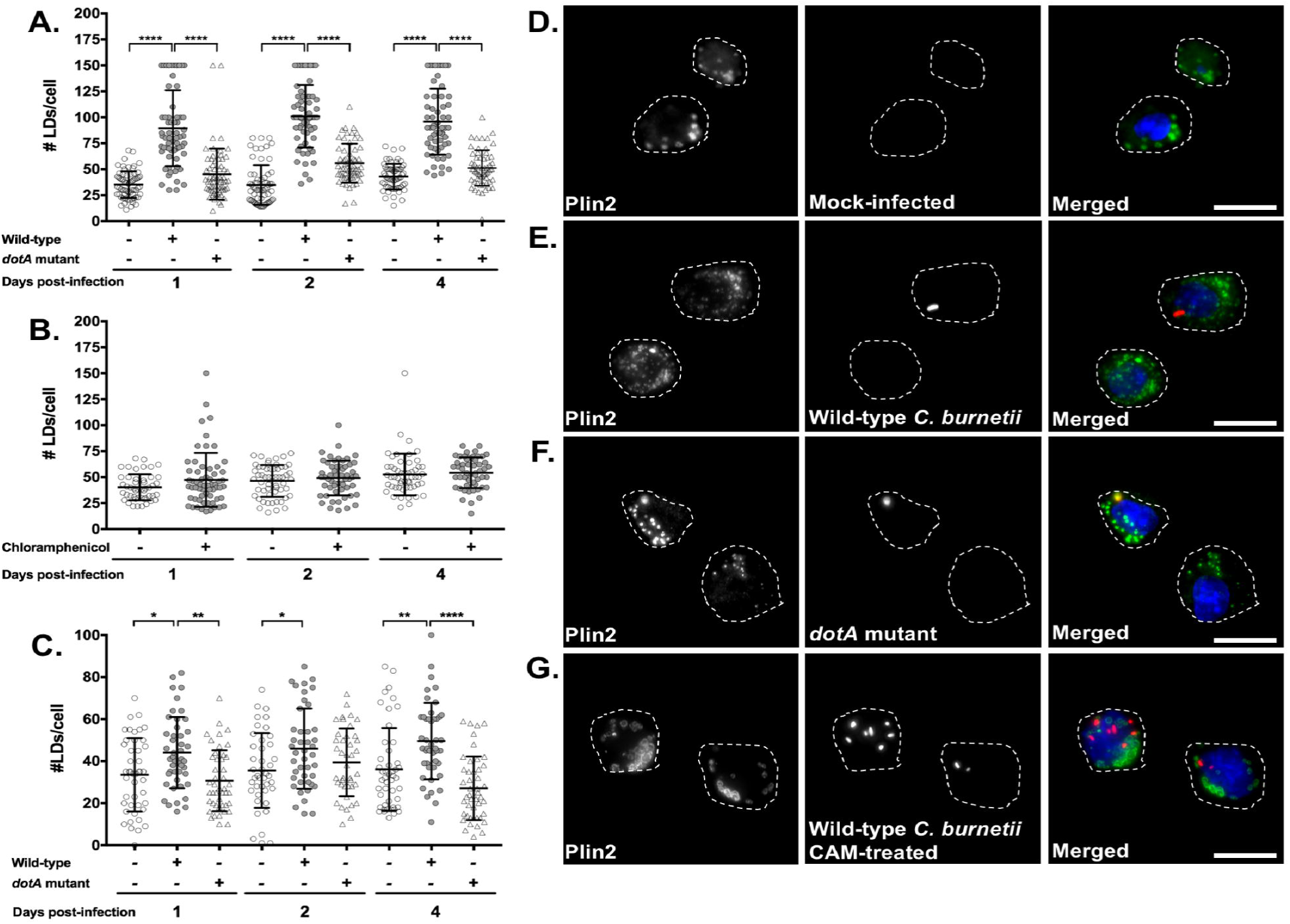
*C. burnetii* infection leads to host cell LD accumulation and is dependent on bacterial protein synthesis and the T4BSS. Macrophages were infected with C. *burnetii* and at different times post-infection, cells were stained for PLIN2 (LDs; green), C. *burnetii* (red) and nucleus (blue). LD number per cell were quantitated by fluorescence microscopy. A) LD numbers in wild-type C. *burnetii* and *dotA* mutant-infected MH-S macrophages. B) LD numbers in MH-S macrophages infected with wild-type C. *burnetii* and treated with chloramphenicol (3ug/ml). C) LD numbers in wild-type C. *burnetii* and *dotA* mutant-infected T HP-1 macrophage-like cells. D) Images of LDs and bacteria in MH-S cells at day 1 post-infection imaged at lO0X. Scale bar = 10 μm. Error bars show the mean of 3 independent experiments +/- SEM * =p<0.05, ** =p<0.01, *** =p <0.001 as determined by ordinary one-way ANOVA with Tukey post-hoc test. Wild-type C. *burnetii* growth in infected MH-S cells treated with different inhibitors was measured at 2 and 4 days post-infection by FFU assay. A-E) Representative images for wild-type MH-S macrophages treated with inhibitors, fixed, stained for PLIN2 (LDs; green) and C. *burnetii* (red) and imaged day 4 post-treatment at lO0X. Scale bar = 10 μm. F) Growth while inhibiting LD formation with triacsin C (10 μM) in wild-type MH-S macrophages. Error bars represent the mean of 4 independent experiments +/- SEM. **= p <0.01 compared to vehicle-treated cells as determined by two-way ANOVA with Bonferroni post-hoc test. G) ACATl protein expression in wild-type and *acat-J 1* macrophages. Cell lysates were immunoblotted and ACATl protein levels were compared with GAPDH as loading control. H) C. *burnetii* growth in vehicle-treated wild-type and *acat-1 1* MH-S macrophages and (I) T863-treated *acat-J-1* MH-S macrophages. Error bars represent the mean of at least 3 independent experiments +/SEM., * =p<0.05, *** =p <0.001 as determined by two-way ANOVA with Bonferroni post-hoc test.

### Host cell LD accumulation is dependent on the *C. burnetii* Type 4B Secretion System (T4BSS)

*C. burnetii* manipulates the host cell by secreting bacterial effector proteins through the T4BSS into the host cytoplasm. To further decipher *C. burnetii’s* role in LD accumulation, we analyzed LD numbers at 1, 2, and 4 days after infection with a *C. burnetii* T4BSS *dotA* mutant (37). While the wild-type *C. burnetii*-infected cells had more than 90 LDs per cell (Figure 2A, closed circles; Figure 2E), T4BSS-mutant infected cells contained an average of 50 LDs, similar to mock-infected cells (Figure 2A, triangles; Figure 2F). This suggests the LD accumulation in *C. burnetii*-infected alveolar macrophages involves the *C. burnetii* T4BSS.

To confirm the T4BSS-dependent increase in LD accumulation, we analyzed LD numbers in human macrophage-like cells (THP-1). When compared to mock- or T4BSS mutant-infected cells, wild-type *C. burnetii*-infected THP-1 cells had increased LD numbers at 1 and 4 days post-infection (Figure 2C), similar to mouse alveolar macrophages. Interestingly, we did not observe T4BSS-dependent LD accumulation at 2 days post-infection. Overall these results demonstrate that *C. burnetii* induces LD accumulation in human and mouse macrophages through a T4BSS-dependent process.

### Blocking LD formation increases *C. burnetii* growth

Given our finding that *C. burnetii* appears to actively manipulate host LDs through the T4BSS, we next assessed the importance of LDs during *C. burnetii* infection. We first blocked LD formation using triacsin C, a long chain fatty acyl CoA synthetase inhibitor (38). Compared to vehicle control, triacsin C significantly reduced macrophage LDs, with <5 LDs per cell (Figure 3A-B). We next treated macrophages with triacsin C during *C. burnetii* infection, and quantitated bacterial growth using a fluorescent infectious focus-forming unit (FFU) assay. At various times post-infection, we recovered bacteria from MH-S cells, replated bacteria onto a monolayer of Vero cells, and incubated for 5 days. After staining for *C. burnetii*, we counted the number of fluorescent foci, with 1 focus unit equivalent to 1 viable bacterium. Surprisingly, compared to vehicle-treated cells, triacsin C treatment increased *C. burnetii* growth 5-fold at 4 days post-infection (Figure 3F).

**Figure 3:**
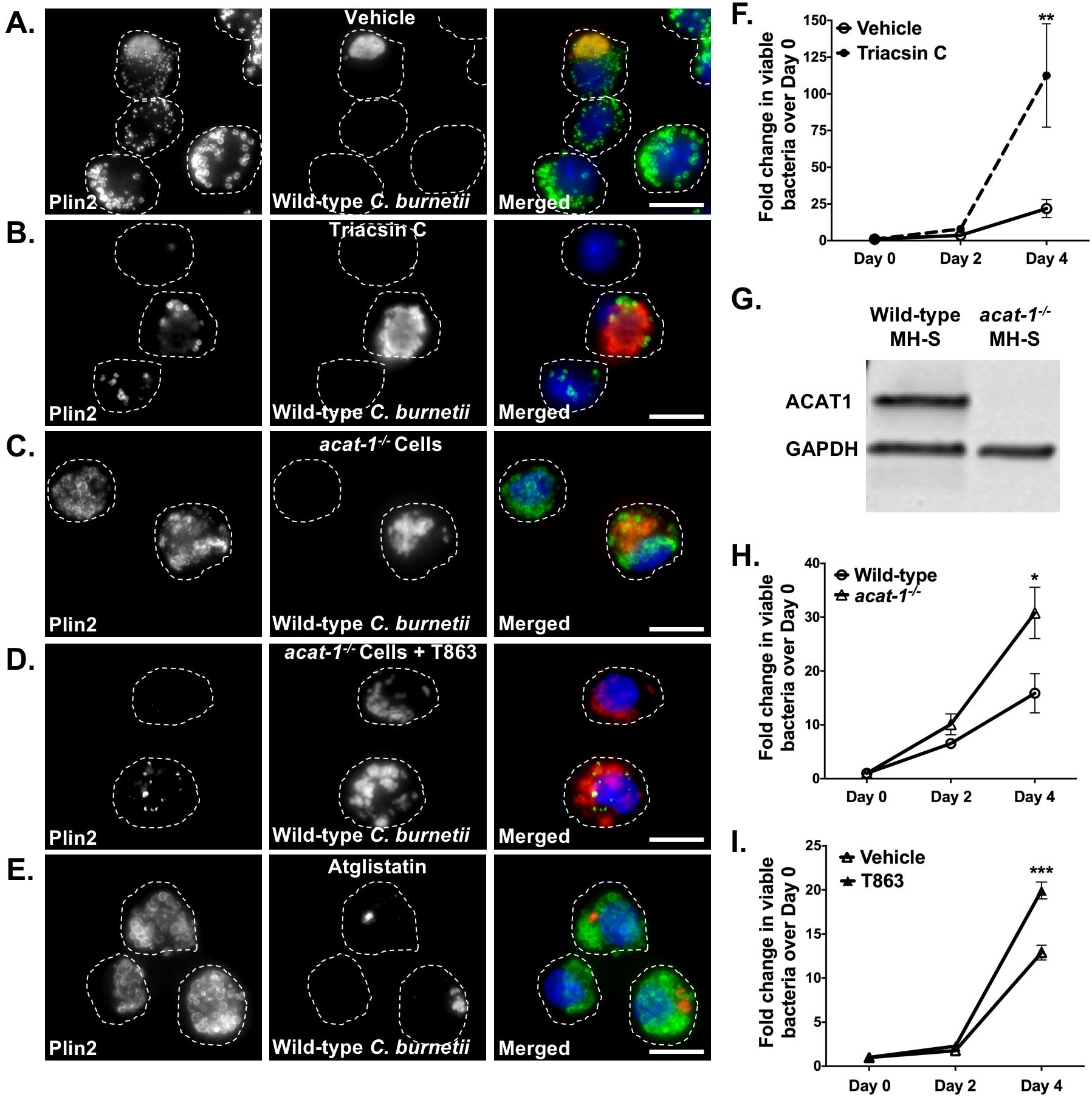
Blocking LD formation increases *C.* ***burnetii*** growth. Wild-type C. *burnetii* growth in infected MH-S cells treated with different inhibitors was measured at 2 and 4 days post-infection by FFU assay. A-E) Representative images for wild-type MH-S macrophages treated with inhibitors, fixed, stained for PLIN2 (LDs; green) and C. *burnetii* (red) and imaged day 4 post-treatment at lO0X. Scale bar = 10 μm. F) Growth while inhibiting LD formation with triacsin C (10 μM) in wild-type MH-S macrophages. Error bars represent the mean of 4 independent experiments +/- SEM. **= p <0.01 compared to vehicle-treated cells as determined by two-way ANOVA with Bonferroni post-hoc test. G) ACATl protein expression in wild-type and *acat-J 1* macrophages. Cell lysates were immunoblotted and ACATl protein levels were compared with GAPDH as loading control. H) C. *burnetii* growth in vehicle-treated wild-type and *acat-1 1* MH-S macrophages and (I) T863-treated *acat-J-1* MH-S macrophages. Error bars represent the mean of at least 3 independent experiments +/SEM., * =p<0.05, *** =p <0.001 as determined by two-way ANOVA with Bonferroni post-hoc test.

To further confirm this finding, we used CRISPR/Cas-9 to knockout *acat-1* (Figure 3G), a gene functionally responsible for sterol esterification in macrophages (39) and upregulated in *C. burnetii*-infected cells (Figure 1). While *acat-1^−/-^* LDs lack sterol esters, fluorescence microscopy revealed similar number of LDs in wild-type and *acat-1^−/-^* cells (Figure 3A,C). Compared to wild-type cells, *C. burnetii* growth in *acat-1^−/-^* cells increased 2-fold at 4 days post-infection (Figure 3H), indicating that blocking sterol esterification favors *C. burnetii* growth. To further deplete both TAG- and sterol ester-containing LDs, we treated *acat-1^−/-^* cells with the DGAT-1 inhibitor T863, which specifically blocks formation of TAGs (40). T863 treatment significantly reduced LDs in *acat-1^−/-^* macrophages, compared to untreated wild-type or *acat-1^−/-^* macrophages (Figure 3A, C, D). *C. burnetii* growth increased 2-fold in T863-treated *acat-1^−/-^* cells compared to vehicle-treated *acat-1^−/-^* cells (Figure 3I), demonstrating that blocking both TAG- and sterol ester-containing LDs improves *C. burnetii* growth.

### Inhibiting LD breakdown blocks *C. burnetii* growth

Because blocking LD formation appeared to benefit *C. burnetii,* we next examined *C. burnetii* growth after inhibiting LD breakdown. When cells or tissues require fatty acids or cholesterol, cytosolic lipases, including adipose triglyceride lipase (ATGL) (3) and hormone sensitive lipase (HSL) (2, 4), hydrolyze TAGs and sterol esters stored in LDs. ATGL catalyzes the initial step of lipolysis converting TAGs to DAGs whereas HSL hydrolyses DAGs to MAGs (41). To block LD breakdown, we first inhibited HSL with a selective inhibitor CAY10499 (42). HSL inhibitor-treated cells showed no significant difference in *C. burnetii* growth compared to vehicle-treated cells (Figure 4A) suggesting conversion of DAGs to MAGs is not important for bacterial intracellular growth.

**Figure 4:**
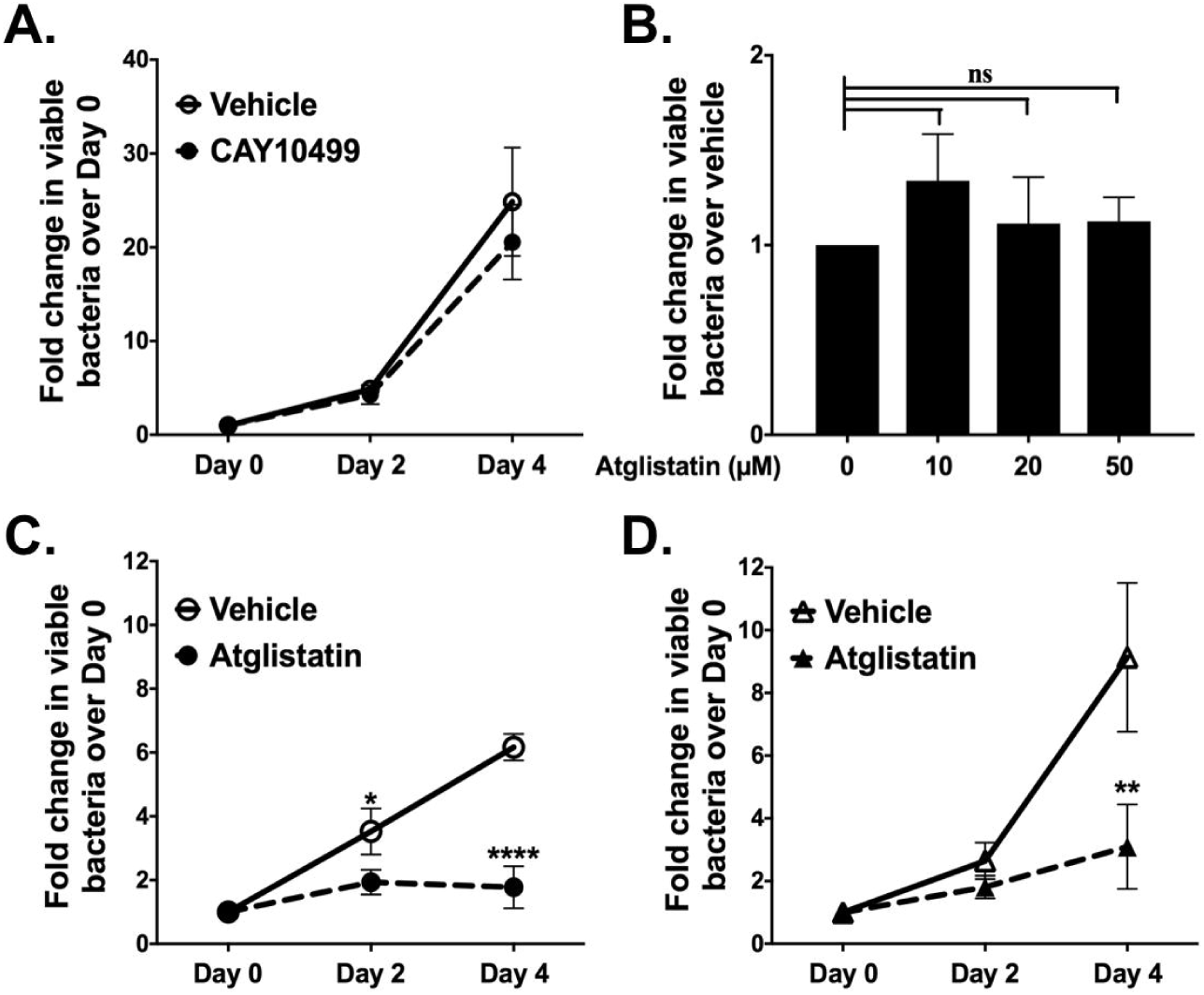
Inhibiting LD breakdown blocks *C. burnetii* growth. Effect of HSL inhibitor CAY10499 and ATGL inhibitor atglistatin on viability of axenic and intracellular wild-type C. *burnetii* was determined by FFU assay. A) C. *burnetii* growth in CAY10499-treated (10 μM) and vehicle-treated wild type MH-S macrophages. Error bars represent the mean of 3 independent experiments +/- SEM. ns = not significant compared to vehicle-treated cells as determined by two-way ANOVA with Bonferroni post-hoc test. B) Direct effect of atglistatin on wild-type C. *burnetii.* Atglistatin was added to axenic C. *burnetii* cultures and bacterial viability was determined at day 4 using FFU assay. Error bars represent the mean of 3 independent experiments +/- SEM. ns = not significant compared to vehicle treatment as determined by ordinary oneway ANOVA with Tukey post-hoc test. C) C. *burnetii* growth in atglistatin-treated (20 μM) and vehicle-treated wild-type MH-S macrophages. Error bars represent the mean of 4 independent experiments +/- SEM. *= p <0.05, ****= p <0.0001 compared to vehicle-treated cells as determined by two-way ANOVA with Bonferroni post-hoc test. D) C. *burnetii* growth in atglistatin-treated (20 μM) and vehicle-treated *acat −1^−/-^*MH-S macrophages. Error bars represent the mean of 4 independent experiments +/- SEM. **= p <0.01 compared to vehicle-treated cells as determined by two way ANOVA with Bonferroni post-hoc test.

To further study the importance of LD breakdown, we inhibited ATGL with the selective and competitive inhibitor atglistatin, which binds the ATGL patatin-like phospholipase domain (43). To eliminate the possibility of ATGL inhibiting a *C. burnetii* phospholipase, we first measured viability of axenic *C. burnetii* cultures in the presence or absence of atglistatin (Figure 4B). Treatment for 4 days had no effect on axenic bacterial growth, indicating atglistatin does not directly affect *C. burnetii*.

We next tested the effect of atglistatin on intracellular bacteria. After atglistatin treatment of wild-type MH-S cells, we observed larger LDs, although the number did not significantly increase (Figure 3E). Interestingly, *C. burnetii* intracellular growth in atglistatin-treated wild-type MH-S cells decreased 5-fold, with essentially no growth (Figure 4C). Further, atglistatin-treated *acat-1^−/-^* cells, which contain TAG-rich LDs, also showed reduced bacterial growth (Figure 4D) suggesting TAG hydrolysis to DAG is critical to support *C. burnetii* growth. Together, these data demonstrate that blocking LD breakdown, particularly TAG to DAG conversion significantly inhibits intracellular *C. burnetii* growth.

## DISCUSSION

Several intracellular pathogens intercept host cell LDs to promote their growth. While certain viruses utilize LDs as platforms for assembly, many bacteria and parasites facilitate intracellular growth by exploiting LDs for nutrition. To understand the role of LDs during *C. burnetii* infection, we assessed changes in LD numbers in *C. burnetii*-infected cells and determined the effect of manipulating LD homeostasis on bacterial growth. Our studies revealed a bacterial T4BSS-dependent increase in LD numbers in *C. burnetii*-infected macrophages. Further, manipulating LD homeostasis significantly altered *C. burnetii* growth. While blocking LD formation promoted *C. burnetii* growth, inhibiting LD breakdown, particularly hydrolysis of TAGs to DAGs, dramatically decreased bacterial growth. Collectively, our data strongly suggest LDs play an important role during *C. burnetii* intracellular growth.

LD formation requires sterol and fatty acid esterification by the enzymes ACAT and DGAT, while LD lipolysis is catalyzed by HSL and ATGL. Once formed, LD maintenance requires the LD-associated proteins PLINs. Previously, two separate microarray analyses reported upregulation of *plin-2* in *C. burnetii*-infected human macrophage-like cells (THP-1) (29, 30), suggesting *C. burnetii* induced changes in host cell LDs. Although we did not observe changes in *plin-2* transcript levels, post-translational regulation of *plin-2* (44) could potentially contribute to changes in LD numbers in infected macrophages. However, consistent with the upregulation of LD-formation genes *acat* and *fabp-4,* our results demonstrated that LD numbers increase in *C. burnetii*-infected mouse and human macrophages.

Notably, LD accumulation occurred as early as day 1 post-infection, when the PV has not expanded and the bacteria are not in log growth. This suggests that LD accumulation is not a host response to a large and growing PV but could be in response to bacterial ligands or due to a process actively manipulated by *C. burnetii* itself. Previously, based on microarray analysis of *C. burnetii*-infected cells where bacterial protein synthesis was blocked, Mahapatra *et al.* identified 36 host cell genes specifically regulated by *C. burnetii* proteins during early stages of infection. These genes were predominantly involved in the innate immune response, cell death and proliferation, vesicular trafficking, cytoskeletal organization and lipid homeostasis. Interestingly, changes in *plin-2* expression level in infected cells was dependent on *C. burnetii* protein synthesis (29) suggesting a role for bacterial proteins in LD accumulation. In agreement with this data, we found that blocking *C. burnetii* protein synthesis prevented increased LD accumulation in mouse alveolar macrophages at all analyzed times post-infection. These data together suggest that *C. burnetii* actively manipulates host LDs at early stages of infection.

Beginning 1 hour post-infection in bone marrow-derived macrophages and 8 hours in HeLa cells, *C. burnetii* secretes effector proteins via T4BSS (45) to modulate host cell functions and promote its intracellular survival. Intriguingly, a *C. burnetii* T4BSS mutant did not increase host cell LD numbers, suggesting a T4BSS-dependent increase in LD accumulation during wild-type *C. burnetii* infection. The same result was observed in both mouse and human macrophages, demonstrating that the *C. burnetii*-induced increase in LDs is independent of species and dependent on the *C. burnetii* T4BSS. This finding suggests that one or more T4BSS effector proteins may actively manipulate LDs, possibly by directly targeting proteins involved in LD homeostasis. For example, the *Salmonella* Typhimurium type 3 secretion system (T3SS) effector protein SseJ esterifies cholesterol and increases LD numbers when ectopically expressed in epithelial and macrophage cells (20). The *C. trachomatis* secreted protein Lda3 localizes to the LD surface and is involved in LD translocation into the *Chlamydia*-containing inclusion (22). Thus far, none of the identified *C. burnetii* T4BSS secreted effector proteins localize to host LDs or associate with proteins involved in LD homeostasis. It is also possible that rather than *de novo* formation, increased LD numbers result from fission of preexisting LDs (46). As observed by microscopy, the LD size in *C. burnetii*-infected macrophages appeared smaller than the mock- or T4BSS mutant-infected macrophages. Smaller, more numerous LDs might result from *C. burnetii* T4BSS-mediated fission of the existing LDs.

While *C. burnetii* T4BSS effector proteins might directly target LD homeostasis pathways, LD accumulation may also be a host innate immune response. In other diseases, LD accumulation occurs during the inflammatory response in macrophages in atherosclerotic lesions (47), leukocytes from joints of patients with inflammatory arthritis (48), and eosinophils in allergic inflammation (49). Thus, an innate immune response to the T4BSS apparatus or T4BSS effector proteins may increase LD numbers in *C. burnetii-*infected macrophages. Additionally, bystander response to bacterial components including TLR ligand LPS could result in increased LD accumulation (16, 50). However, T4BSS mutant and protein-synthesis blocked *C. burnetii* have similar surface ligands to wild-type bacterium but show no increase in LD accumulation thus arguing against the contribution of *C. burnetii* LPS and other surface ligands. Overall, while our data demonstrate that the *C. burnetii* T4BSS is involved in LD accumulation in both mouse and human macrophages, the bacterial effector proteins and the specific LD processes targeted remain unknown.

Besides T4BSS-dependent increase in LD accumulation, we found that manipulating LD homeostasis, in particular fatty acids, significantly altered *C. burnetii* growth. Surprisingly, blocking LD formation by pharmaceutical and genetic approaches, which increases the availability of free fatty acids, led to improved *C. burnetii* fitness in macrophages. Conversely, reducing fatty acid availability by inhibiting breakdown of LD-derived TAGs to DAGs blocked *C. burnetii* growth. Intriguingly, DAG to MAG conversion did not affect bacterial growth, suggesting that DAGs in particular are critical for *C*. *burnetii* infection. DAGs and other fatty acids liberated during LD breakdown can be re-esterified or serve as signaling cofactors, building blocks for membranes, or substrates for β-oxidation (2). Other bacterial pathogens are known to target LDs as a source of lipids. For example, *M*. *tuberculosis* uses free fatty acids as a source of energy and carbon (15, 51), while *C. trachomatis* is hypothesized to break down bacterial inclusion-associated host LDs to provide lipids for bacterial growth (14). It not known if free fatty acids or sterols liberated from LDs, either in the cytosol or possibly in PV lumen, are used directly by *C. burnetii*.

In addition to serving as a source of free fatty acids and sterols, macrophage LDs are rich in substrates and enzymes that generate prostaglandins and leukotrienes, which are arachidonic acid-derived inflammatory lipid mediators (52, 53). In *M. leprae*-infected Schwann cells and *M. bovis* BCG-infected macrophages, increased LD biogenesis correlates with increased production of prostaglandin E2 (PGE2), linking LDs to the production of innate immune modulators (17, 54). PGE2 has a potential role in inhibiting TH1 responses important in clearance of intracellular pathogens (24) including *C. burnetii* (55). Interestingly, elevated levels of PGE2 were observed in *C. burnetii* endocarditis patients and linked to *C. burnetii*-mediated immunosuppression. Koster *et al.* reported lymphocytes from chronic Q fever patients being unresponsive to *C.burnetii* antigens, an effect reversed by PGE2 suppression with indomethacin (56). In addition, after stimulation with *C. burnetii* antigens, monocytes from Q fever patients produced PGE2, which in turn downregulated T lymphocyte-mediated IL-2 and IFNγ production. Interestingly, PGE2 synthesis inhibitor Piroxicam reversed this downregulation of pro-inflammatory cytokine production (57). Thus, while PGE2 appears to play a role in Q fever patients, the relationship between *C. burnetii*-induced LDs and PGE2 production is not known. Considering that LD breakdown can serve multiple functions, *C. burnetii* could use LDs either as a source of nutrients or for production of lipid immune mediators like PGE2, which could then modulate the host cell response to promote *C. burnetii* intracellular growth.

In summary, our data demonstrate that LD homeostasis is important for *C. burnetii* intracellular survival. Because the *C. burnetii* T4BSS is involved in LD accumulation, characterizing bacterial T4BSS effector proteins that target host LD homeostasis will help further understand the role of LDs in *C. burnetii* pathogenesis.

## MATERIALS AND METHODS

### Bacteria and mammalian cells

*C. burnetii* Nine Mile Phase II (NMII; clone 4, RSA439) were purified from Vero cells (African green monkey kidney epithelial cells, ATCC CCL-81; American Type Culture Collection, Manassas, VA) and stored as previously described (58). For experiments examining T4BSS-dependent accumulation of LDs, NMII and the *dotA* mutant (37) were grown for 4 days in ACCM-2, washed twice with phosphate buffered saline (PBS) and stored as previously described (59). Vero, mouse alveolar macrophages (MH-S; ATCC CRL-2019) and human monocytes (THP-1; ATCC TIB-202) were maintained in RPMI (Roswell Park Memorial Institute) 1640 medium (Corning, New York, NY, USA) containing 10% fetal bovine serum (Atlanta Biologicals, Norcross, GA, USA) at 37°C and 5% CO_2_ and human embryonic kidney 293 (HEK293T; ATCC CRL-3216) in DMEM (Dulbecco’s Modified Eagle Medium) (Corning, New York, NY, USA) containing 10% fetal bovine serum at 37°C and 5% CO_2_. THP-1 cells were differentiated with 200 nM of phorbol 12-myristate 13-acetate (PMA) for 24 hours. PMA was removed, and the cells rested for 48 hours prior to infection. The multiplicity of infection (MOI) was optimized at 37^°^C and 5% CO_2_ for each bacterial stock, cell type and infection condition for a final infection of ∼1 internalized bacterium/cell.

### Generating *acat-1^−/-^* MH-S cell line

The guide RNA sequence 5‘TCGCGTCTCCATGGCTGCCC3’ to mouse *acat-1* was selected using the optimized CRISPR design site crispr.mit.edu. Oligonucleotides were synthesized (IDT, Coralville, IA, USA), annealed, and cloned into the lentiCRISPRv2 plasmid (a gift from Feng Zhang, Addgene # 52961, Cambridge, MA, USA) (60), at the BsmBI restriction site to generate plentiCRISPRv2-*acat-1*. To generate lentivirus, HEK293T cells were co-transfected with plentiCRISPRv2-*acat-1* and packaging plasmids pVSVg (Addgene # 8454), pRSV-Rev (Addgene # 12253), and pMDLg/pRRE (Addgene # 12251) using FuGENE6 reagent (Promega, Madison, WI, USA). At 48 hours post-transfection, supernatant was collected, centrifuged at 3000xg, and then filtered with 0.45μm filter to remove cells and debris. Supernatant was concentrated using the Lenti-X concentrator (Catalog # PT4421-2, Clontech, USA) and viral RNA isolated using Viral RNA isolation kit (Catalog # 740956, Macherey-Nagel, Germany) to determine viral titer using Lenti-X qRT-PCR titration kit (Catalog # PT4006-2, Clontech). Viral titers were optimized for transduction of MH-S cells to generate stable *acat-1^−/-^* cells.

2x10^5^ MH-S cells were plated in a 6 well plate and transduced with 5.8 x 10^6^ viral particles/ml. 1μg/ml puromycin was used for selection 48 hours post-transduction and continued for 24 hours. The puromycin was then removed and the cells allowed to recover before isolating individual clones by limiting dilution.

To confirm disruption of *acat-1*, clones were lysed in 2% SDS (Sigma-Aldrich, St. Louis, MO, USA) for SDS-PAGE and immunoblotting with 1:1000 rabbit anti-mouse ACAT1-specific antibody (Catalog # NBP189285, Novus Biologicals, Littleton, CO, USA) and 1:4000 GAPDH loading control monoclonal antibody (Catalog # MA5-15738, ThermoFisher Scientific, Waltham, MA, USA).

### Gene expression analysis of LD genes

2x10^5^ MH-S cells were infected in a 6 well plate. At day 1 post-infection, host cell RNA was isolated using RNEasy mini kit (Catalog # 74104, Qiagen) and reverse transcribed to cDNA using Superscript first strand synthesis system (Catalog # 11904-018, Invitrogen). Gene expression analysis was performed using TaqMan array mouse lipid regulated genes (Catalog # 4415461, Applied Biosystems). Fold change was calculated compared to mock-infected cells in 6 independent experiments.

### LD quantitation

1x10^5^ MH-S cells were plated onto ibidi-treated channel μslide VI^0.4^ (3x10^3^ cells per channel; Ibidi, Verona, WI) and allowed to adhere overnight. After infecting with *C. burnetii* for 1 hour, cells were gently washed with phosphate buffered saline (PBS) to remove extracellular bacteria, and incubated in 10% FBS-RPMI. For blocking bacterial protein synthesis, infected cells were incubated with 3 μg/ml chloramphenicol containing 10% FBS-RPMI for the remainder of experiment, replenishing media every 24 hours. At different times post-infection, infected cells were fixed with 2.5% paraformaldehyde on ice for 15 min, then permeabilized/blocked for 15 min with 0.1% saponin and 1% bovine serum albumin (BSA) in PBS (saponin-BSA-PBS) and stained with 1:1000 rabbit anti-mouse PLIN2 primary antibody (Catalog # PA1-16972, ThermoFisher Scientific), 1:2000 guinea-pig anti-*C. burnetii* primary antibody (61) and 1:1000 rat anti-LAMP (Catalog # 553792, BD Biosciences) primary antibody in saponin-BSA-PBS for 1 hour. THP-1 cells were stained with 1:500 guinea-pig anti-human PLIN2 primary antibody (Catalog # 20R-AP002, Fitzgerald Industries International, Acton, MA), 1:2000 rabbit anti-*C. burnetii* primary antibody and 1:1000 rat anti-LAMP primary antibody in saponin-BSA-PBS for 1 hour. After three washes with PBS, cells were stained with 1:2000 AlexaFluor 488 anti-rabbit, AlexaFluor 594 anti-guinea pig and AlexaFluor 647 anti-rat secondary antibodies (Invitrogen) for 1 hour. ProLong Gold mount (Invitrogen) was added to the wells after washing with PBS and slides visualized on a Leica inverted DMI6000B microscope (100X oil). The number of LDs per cell was quantitated for 25 cells per condition in duplicate in each of three individual experiments, with only bacteria-containing cells counted for *C. burnetii*-infected cells.

### Inhibitors

Each LD homeostasis inhibitor used was diluted in DMSO based on manufacturer’s instructions and optimum inhibitor concentration was determined based on 100% host cell viability determined by trypan blue staining, and changes in LD numbers per cell. The optimum concentrations determined for each inhibitor was: Triacsin C (Enzo Life Sciences, Farmingdale, NY, USA) – 10 μM, T863 (Sigma-Aldrich) – 10 μM, CAY10499 (Cayman Chemicals, Ann Arbor, MI, USA) - 10 μM, Atglistatin (Cayman Chemicals) – 20 μM.

### *C. burnetii* growth by fluorescent infectious focus-forming unit (FFU) assay

To measure growth of *C. burnetii* in wild-type and *acat-1^−/-^* MH-S cells, 5x10^4^ cells/well were infected for 1 hour in a 48 well plate, washed with PBS, and then incubated with media containing respective vehicle and inhibitors. At the indicated time points, the media was removed and cells were incubated with sterile water for 5 min, pipetted up and down and the lysate diluted 1:5 in 2% FBS-RPMI. Serial dilutions were added to 24 well plate containing confluent monolayers of Vero cells, incubated for 5 days, fixed with methanol and stained with rabbit anti-*C. burnetii* antibody as well as DAPI to confirm monolayer integrity. Four fields per well were captured on an Evos automated microscope (ThermoFisher) with 4X objective and fluorescent foci units were quantitated using ImageJ. Each experiment was done in duplicate.

### Atglistatin-treatment of *C. burnetii* axenic cultures

To test bacterial sensitivity to atglistatin, ACCM-2 was inoculated at approximately 1x10^5^ bacteria/ml with *C. burnetii* NMII and grown for 3 days as previously described (59). Bacteria (500 μl) were then incubated with DMSO or atglistatin in 24 well plates under normal *C. burnetii* culture conditions. Media was replenished every 24 hours by centrifuging the supernatant at 20000xg for 10 min, and bacterial pellet resuspended in new media containing inhibitor. After 4 days, bacteria were diluted 1:10 in 2% FBS-RPMI and serial dilutions were added to confluent Vero cell monolayers in a 96 well ibidi-treated μplate. At 5 days post-infection, the plate was stained and fluorescent foci were determined as above. Each experiment was done in duplicate.

### Statistical analysis

Statistical analyses were performed using two-tailed paired t-test, ordinary one-way ANOVA or two-way ANOVA with Tukey’s or Bonferroni’s multiple comparisons test in Prism (GraphPad Software, Inc., La Jolla, CA).

## ACKNOWLEDGMENTS

This research was supported by National Institutes of Health (AI21786; S.D.G.), American Heart Association (16POST27250157; M.M.) and National Institutes of Health (5R25GM07592; B.Z.). We thank Anna Justis, Tatiana Clemente and Rajshekar Gaji for critical reading of the manuscript and members of the IU Biology of Intracellular Pathogens Group for helpful suggestions.

We have no conflicts of interest to declare.

